# Systems level phosphoproteomics reveals CAMKK2 driven kinase signaling underlying malignant phenotypes in gastric cancer

**DOI:** 10.64898/2026.02.06.704339

**Authors:** Mohd. Altaf Najar, T. S. Keshava Prasad, Prashant Kumar Modi

## Abstract

Gastric cancer is driven by aberrant kinase signaling that promotes uncontrolled proliferation and malignant progression. Calcium/calmodulin-dependent protein kinase kinase 2 (CAMKK2) is overexpressed in gastric cancer; however, the global phosphorylation networks downstream of CAMKK2 remain incompletely defined. In this study, we investigated the functional and signaling consequences of CAMKK2 inhibition in gastric cancer cells using an integrated phenotypic and quantitative phosphoproteomics approach. Pharmacological inhibition of CAMKK2 using STO-609 in AGS cells significantly suppressed proliferation, clonogenic growth, migration, and invasion, and induced defects in nuclear morphology indicative of impaired cell cycle progression. Tandem mass tag (TMT) based phosphoproteomic profiling identified over 10,500 phosphopeptides and revealed extensive phosphoproteome remodeling following CAMKK2 inhibition, characterized predominantly by hypophosphorylation of proteins involved in nuclear signaling, RNA processing, and cell cycle regulation. Kinase substrate enrichment and motif analyses demonstrated coordinated attenuation of CDK, MAPK, and mitotic kinase-associated signaling pathways, with convergence on E2F regulated transcriptional programs. Collectively, these findings establish CAMKK2 as a central regulator of kinase signaling networks that sustain proliferative and malignant phenotypes in gastric cancer and highlight CAMKK2 inhibition as a potential therapeutic strategy.

## Introduction

Gastric cancer (GC) remains one of the most lethal malignancies worldwide, with particularly high incidence and mortality among older males [1]. The global burden of GC varies markedly across geographical regions and is strongly influenced by dietary habits and *Helicobacter pylori* infection [1, 2]. According to GLOBOCAN 2020, gastric cancer ranks as the fifth most diagnosed cancer and the fourth leading cause of cancer-related mortality globally [3]. Despite advances in diagnosis and treatment, the molecular mechanisms driving gastric tumorigenesis and disease progression remain incompletely understood, limiting the development of effective targeted therapies.

At the molecular level, gastric cancer is characterized by the dysregulation of multiple signaling pathways resulting from activation of proto-oncogenes and inactivation of tumor suppressor genes [4]. Aberrant regulation of the cell cycle is a central feature of gastric carcinogenesis, with genetic and signaling alterations converging on key cell cycle regulators. Proteins governing the G1-S phase transition, including Cyclin D1 and the retinoblastoma-associated protein (RB1), play critical roles in controlling cell proliferation and are frequently deregulated in GC [5]. Cyclin D1 overexpression has been reported across multiple stages of gastric cancer, from early lesions to advanced disease, underscoring the importance of kinase-driven cell cycle control in GC pathogenesis.

Protein kinases are central mediators of intracellular signal transduction and play essential roles in regulating cell growth, proliferation, survival, and differentiation. Dysregulation of kinase signaling is a hallmark of cancer and is frequently associated with uncontrolled cellular proliferation and tumor progression [6]. In gastric cancer, aberrant activation of multiple kinase-driven pathways, including EGFR, HER2, and Ras/MAPK signaling, has been extensively documented [4]. Several kinases have been reported to be activated or deregulated in GC, including mitogen-activated protein kinases ERK1 and ERK2, KRAS, RAF1, cyclin-dependent kinase 5 (CDK5), and calcium/calmodulin-dependent protein kinase kinase 2 (CAMKK2) [7–10]. CAMKK2 is a 66–68 kDa serine/threonine kinase composed of unique N- and C-terminal domains and a central kinase domain followed by a regulatory region containing overlapping autoinhibitory and calmodulin-binding motifs [11, 12]. As a calcium/calmodulin-responsive kinase, CAMKK2 functions as an upstream regulator of multiple signaling cascades and is known to phosphorylate and activate downstream kinases such as CAMKI, CAMKIV, and AMPK [12–14]. Serine/threonine kinases, including CAMKK2, are critical for maintaining cellular homeostasis through regulation of transcription factors, metabolic enzymes, and cell cycle regulators, and their dysregulation has been implicated in tumor growth and metastasis [11, 15].

Accumulating evidence suggests that CAMKK2 is overexpressed in multiple cancer types, including gastric cancer, and contributes to oncogenic phenotypes [8]. CAMKK2 has emerged as a potential therapeutic target in several malignancies, including gastric cancer [16, 17]. However, the downstream signaling mechanisms mediated by CAMKK2 appear to be highly context dependent. In ovarian cancer, CAMKK2 has been reported to promote AKT-mediated signaling [18], whereas in prostate cancer, CAMKK2 signaling is linked to AMPK activation, resulting in increased GLUT12 (SLC2A12) expression and enhanced glucose uptake [19]. These findings indicate that CAMKK2 engages distinct signaling networks depending on cellular and tissue context.

Although we and others have demonstrated that CAMKK2 is overexpressed in gastric cancer [8, 17] and that its inhibition leads to reduced oncogenic properties, including diminished proliferation and transformation, and although our previous work identified a CAMKK2–MEK/ERK–CDK–MCM signaling axis driving gastric cancer cell proliferation, a comprehensive systems-level understanding of CAMKK2-dependent phosphorylation networks remains lacking [16, 17]. In particular, the broader kinase signaling architecture, substrate specificity, and integration of CAMKK2 signaling with additional cellular processes beyond canonical MAPK driven proliferation such as RNA processing, transcriptional regulation, cytoskeletal dynamics, and stress responses have not been systematically explored.

To address this gap, we employed a tandem mass tag (TMT)-based quantitative phosphoproteomics approach to systematically characterize CAMKK2-regulated phosphorylation networks in gastric cancer cells. Using pharmacological inhibition of CAMKK2 in AGS gastric cancer cells, we sought to define the global phosphoproteomic landscape, identify downstream kinase signaling pathways, and elucidate the molecular mechanisms by which CAMKK2 sustains malignant phenotypes in gastric cancer.

## Materials and Methods

### Reagents and Chemicals

The human gastric cancer cell lines AGS were obtained from American Type Culture Collection (ATCC, Manassas, VA), USA. Dulbecco‘s Modified Eagle Medium-high glucose ((DMEM, Cat#12100046)), fetal bovine serum (FBS, Cat#10270-106), and 100X antibiotic/antimycotic solution (Cat#15240-062), Trypsin EDTA (Cat#25200-056), were purchased from Gibco, Thermo Fischer Scientific, USA, and phosphate-buffered saline (PBS, Cat#TL101), from HiMedia, India. All plasticware for cell culture was procured from Nunc.

STO-609 (7-Oxo-7H-benzimidazo[2,1-a]benz[de]isoquinoline-3-carboxylic acid – acetic acid), a CAMKK2 inhibitor, was purchased from Santa Cruz Biotechnology, USA. Thiazolyl blue tetrazolium bromide (MTT, Cat#M5655), bisBenzimide H33342 (HOECHST, Cat#B2261), propidium iodide (PI, Cat#P4170), 2‘,7‘-Dichlorofluorescein diacetate (DCFDA, Cat#D6883), Triton X-100 (Cat#T8787), trisodium citrate (Cat#S1804) were procured from Sigma-Aldrich, USA. Dimethyl sulfoxide (DMSO, Cat#196055) was procured from MP Biomedicals, USA. Sodium dodecyl sulfate (SDS, Cat#L3771), sodium pyrophosphate (Cat#S6422), sodium orthovanadate (Cat#S6508), ammonium persulfate (Cat#A3678), brilliant blue-R (Cat#B7920), Glycerol (Cat#24505) from Fisher Scientific, USA. Tris base (Cat#648310), from Merck, USA. Tetramethylethylenediamine (TEMED, Cat#194019), were procured from MP Biomedicals, India.

Pierce™ BCA protein estimation assay kit (Cat# 23225), Pierce™ Peptide estimation assay kit (Cat# 23275), and TMT 10plex kit (Cat# 90110) were procured from ThermoFisher Scientific USA. Anhydrous acetonitrile (Cat#271004), DL-dithiothreitol (Cat#D9779), iodoacetamide (Cat#I6125), trifluoroacetic acid (Cat#302031), HEPES (Cat#H4034), and formic acid (Cat# 5330020050) were procured from Sigma Aldrich, St. Louis, USA. TPCK-treated Trypsin (Cat#LS003741), was purchased from Worthington Biochemical Corporation, USA. Solid-phase extraction disks, C-18 (Cat#66883-U), were procured from EmporeTM, USA. LC-MS grade solvents for liquid chromatography-mass spectrometry analysis such as acetonitrile (Cat#1.00029.2500), methanol (Cat#1.06035.2500), and water (Cat #1.15333.2500) were procured from Merck, USA.

### Cell culture

AGS (CRL-1739) gastric cancer cells were obtained from the American Type Culture Collection (ATCC, Manassas, VA, USA). Cells were maintained in a humidified incubator at 37 °C with 5% CO₂ and cultured in Dulbecco’s Modified Eagle Medium (DMEM) supplemented with 10% fetal bovine serum (FBS) and 1× antibiotic–antimycotic solution.

### Colony formation assays

Colony formation assays were performed as described previously [16]. Briefly, AGS cells were seeded at a density of 1,500 cells per well in 6-well plates. After 12 h, cells were treated with the CAMKK2 inhibitor STO-609 or vehicle control and allowed to grow for 12–14 days. Colonies were fixed with methanol and stained with 1% crystal violet (Sigma-Aldrich, St. Louis, MO, USA). Images were acquired using an inverted light microscope (Carl Zeiss, Germany) at 8× magnification. Colony number and area were quantified using ImageJ software (NIH, USA). All experiments were performed in triplicate and repeated independently three times. Data are presented as mean ± SEM.

### Wound healing assays

Wound healing assays were performed as described previously [16]. AGS cells were grown to confluence, and a uniform scratch was introduced using a sterile pipette tip. Cells were treated with STO-609 or vehicle control and allowed to migrate for up to 36 h. Wound images were captured at 0 and 36 h using an inverted microscope. All experiments were performed in triplicate unless otherwise stated.

### Invasion assays

Cell invasion assays were performed using a Matrigel-coated transwell system (BD Biosciences, San Jose, CA, USA). Polyethylene terephthalate (PET) membranes with 8-µm pore size were used. AGS cells treated with STO-609 or vehicle control were seeded at a density of 2.0 × 10⁴ cells in 500 µL serum-free medium in the upper chamber. The lower chamber contained complete growth medium. Plates were incubated at 37 °C for 48 h. Non-invading cells were removed from the upper surface of the membrane, and invading cells on the lower surface were fixed and stained with crystal violet. Each experiment was performed in duplicate and repeated three times.

### Cell proliferation assay

Cell proliferation was assessed using crystal violet staining. AGS cells (5 × 10⁵ cells per well) were seeded in 6-well plates and treated with 18.5 µM STO-609 for 72 h. Cells were washed with PBS and stained with 3% crystal violet prepared in 50% methanol for 1 h at room temperature. Excess stain was removed by washing, and images were captured using an inverted microscope (Carl Zeiss, Germany) at 8× magnification.

### Sample preparation for phosphoproteomics analysis

#### Protein isolation and digestion

AGS cells (5 × 10⁶ cells per 10-cm dish) were treated with STO-609 or vehicle control for 2 h at ∼70% confluence. Cells were lysed in buffer containing 8 M urea, 75 mM NaCl, 50 mM Tris-HCl (pH 8.0), 1 mM EDTA, and phosphatase inhibitor cocktail (Thermo Fisher Scientific). Lysates were sonicated on ice and centrifuged at 13,000 × g for 30 min. Protein concentration was determined by BCA assay. One milligram of protein per sample was reduced, alkylated, diluted to 1 M urea using 50 mM TEABC, and digested overnight with TPCK-treated trypsin (1:20 enzyme-to-protein ratio). Peptides were desalted using C18 Sep-Pak cartridges and dried prior to TMT labeling

### Tandem mass tag (TMT) labeling

TMT-based quantitative phosphoproteomics was performed using TMT 6-plex reagents (Thermo Fisher Scientific) as described previously [16]. Control samples were labeled with TMT channels 126, 127, and 128, and STO-609-treated samples with channels 129, 130, and 131. Labeling efficiency was confirmed prior to pooling equal amounts of labeled peptides.

### Phospho-enrichment

Phosphopeptides were enriched using Fe-NTA beads (Thermo Scientific, A32992) according to the manufacturer’s instructions. The flow-through was subsequently subjected to TiO₂-based enrichment as described previously [16] to enhance phosphoproteome coverage.

### Peptide fractionation and clean-up

High-pH reversed-phase peptide fractionation was performed using C18 StageTips with 2% TEABC as mobile phase. Peptides were eluted using increasing concentrations of acetonitrile (5–50%). Fractions were dried, desalted using C18 StageTips, and stored at −20 °C until LC-MS/MS analysis.

### LC-MS/MS analysis

Cleaned and dried phosphopeptides were reconstituted in 0.1% formic acid before mass spectrometry analysis. Orbitrap Fusion Tribrid mass spectrometer (Thermo Fischer Scientific, Bremen, Germany) connected to Easy-nLC-1200 nanoflow liquid chromatography system (Thermo Fischer Scientific) was used for data acquisition. The peptides were loaded onto a 2 cm trap column (nanoViper, 3 µm C18 Aq) (Thermo Fisher Scientific). Peptide separation was done using a 15 cm analytical column (nanoViper, 75 µm silica capillary, 2 µm C18 Aq) at a 300 nL/min flow rate. The solvent gradients were set as a linear gradient of 5-35% solvent B (80% acetonitrile in 0.1% formic acid) for 90 min with the total run time for each fraction was 120 minutes. Global MS survey scan was carried out at a scan range of 400-1600 m/z mass range (120,000 mass resolution at 200 m/z) in a data-dependent mode using an Orbitrap mass analyzer.

The maximum injection time was 5 ms. Only peptides with charge states 2-6 were considered for analysis, and the dynamic exclusion rate was set to 30 s. For MS/MS analysis, data was acquired at top speed mode with 3s cycles and subjected to higher collision energy dissociation with 34% normalized collision energy. MS/MS scans were carried out at a range of 100-2000 m/z using Orbitrap mass analyzer at a resolution of 30,000 at 200 m/z. Maximum injection time was 200ms. Internal calibration was carried out using the lock mass option (m/z 445.1200025) from ambient air.

### Database search for peptide, protein, and phosphopeptide identification

The raw files obtained after the data acquisition were searched using Proteome Discoverer software suite version 2.2 (Thermo Fisher Scientific). The MS/MS data were searched against the human protein database (RefSeq 102 database, along with known mass spectrometry contaminants) using SEQUEST and Mascot algorithms. Search parameters included carbamidomethylation of cysteine, TMT at peptide N-terminal and lysine as a static modification. Dynamic modifications included oxidation of methionine, phosphorylation at serine, threonine, tyrosine, and minimum peptide length was 7 amino acids selected with 1 missed cleavage allowed. Mass tolerance was set to 10 ppm at MS level and 0.05 Da for MS/MS, and the false discovery rate was set to 1%.

### Data analysis

The results file from Proteome Discoverer was used for further analysis. The Perseus 80 tool was used to compute fold change (FC), p-value, and q-value. Morpheus tool (Broad Institute;https://software.broadinstitute.org/morpheus/) was used for generating heat maps. The Sankey diagram was generated using an online Sankey generator (http://sankey-diagram-generator.acquireprocure.com/). Proteins were classified and categorized based on Gene Ontology analysis using DAVID 8.6 functional annotation tool (https://david.ncifcrf.gov/summary.jsp) and pathway analysis using Reactome 83 (https://reactome.org/) tools. The proteins identified were also classified based on their function using data from The Human Protein Altas (HPA) (https://www.proteinatlas.org/). The Gene Ontology (GO) analysis was performed using g:Profiler, a web-based server for functional enrichment analysis [20, 21].

### Kinome Map

The kinome map was built using the KinMap tool (http://www.kinhub.org/kinmap/index.html). The list of identified kinases was searched, and relevant kinases are highlighted on the kinome map. Dysregulated kinases (hyper- or hypo-phosphorylated upon CAMKK2 inhibition) are also depicted.

### Kinase enrichment analysis

Kinase enrichment analysis was done using the online eXpression2Kinases (X2K) (http://amp.pharm.mssm.edu/X2K/) tool. Proteins with decreased phosphorylated upon CAMKK2 inhibition were used for input data analysis.

### PTM profiling and motif analysis

High-confidence PTMs were profiled using posttranslational modification–profiling (PTM-Pro) tool with a ptmRS site probability >75% as per the literature. Motif enrichment was done using PTM-Pro 2.0 against HsfSeq-94 with a cut-off of 75%.

### Statistical analysis

All experiments were performed in biological triplicate. Data is represented as mean ± Standard deviation. For statistical comparison, the data were analyzed by Student’s t-test using GraphPad PrismTM software (version 6.1). p < 0.05 was considered statistically significant.

## Results

CAMKK2 has been reported to be overexpressed in gastric cancer and implicated in the regulation of oncogenic signaling pathways [22]. Consistent with these observations, our previous studies have demonstrated that CAMKK2 promotes cellular transformation and proliferation in gastric cancer cells [17], in part through activation of CDK signaling via the MEK–ERK pathway. In addition, CAMKK2 has been shown to regulate multiple cancer-associated signaling pathways, including the AKT pathway [13, 18, 23].

Given the established role of CAMKK2 in sustaining oncogenic signaling, we sought to directly assess the functional consequences of CAMKK2 inhibition on malignant phenotypes in gastric cancer cells. AGS cells were treated with the CAMKK2 inhibitor STO-609, and the effects on cell proliferation, clonogenic growth, migration, invasion, and nuclear morphology were systematically evaluated (Figure 1). CAMKK2 inhibition resulted in a pronounced suppression of proliferative capacity, reduced clonogenic potential, and impaired migratory and invasive behavior. Furthermore, prolonged CAMKK2 inhibition led to a reduction in cell number accompanied by the emergence of multinucleated cells, suggesting defects in cell cycle progression and mitotic regulation.

**Figure 1.**
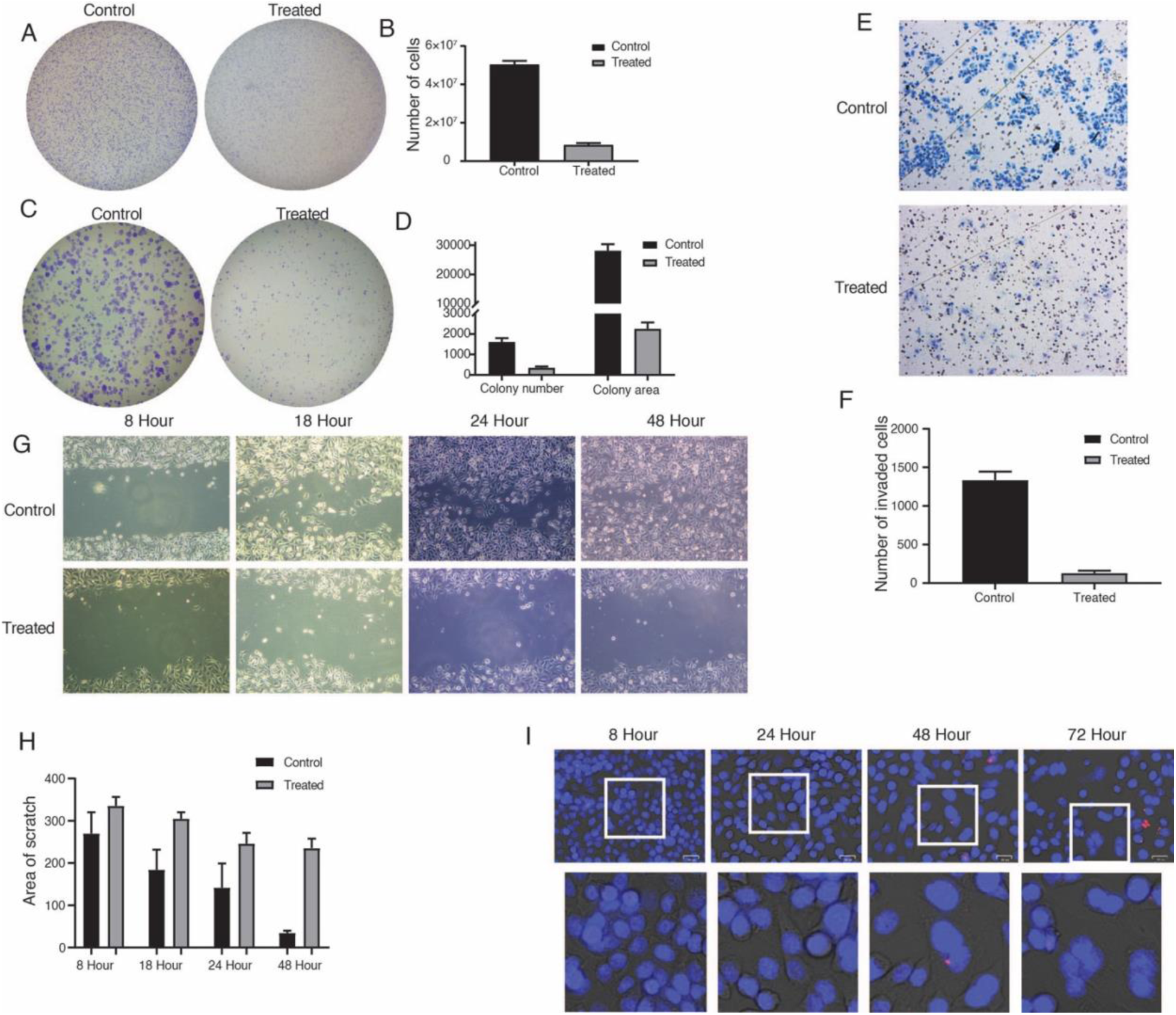
CAMKK2 inhibition suppresses proliferation, clonogenic growth, migration, invasion, and induces nuclear abnormalities in gastric cancer cells: **(A)** Representative images of crystal violet–stained AGS cells showing reduced cell proliferation following treatment with the CAMKK2 inhibitor STO-609 compared to vehicle control. **(B)** Quantification of proliferating cells demonstrating a significant decrease in cell number upon CAMKK2 inhibition. **(C)** Representative images of colony formation assays showing reduced number and size of colonies in STO-609–treated AGS cells compared to control. **(D)** Quantitative analysis of colony number and colony area at indicated time points following CAMKK2 inhibition. **(E)** Representative images from transwell invasion assays showing reduced invasive capacity of AGS cells upon CAMKK2 inhibition. **(F)** Quantification of invaded cells demonstrating significant suppression of invasion following STO-609 treatment. **(G)** Representative wound healing assay images at indicated time points (8, 18, 24, and 48 h) showing impaired migratory capacity in CAMKK2-inhibited cells compared to control. **(H)** Quantitative analysis of wound closure area demonstrating delayed wound healing upon CAMKK2 inhibition. **(I)** Representative DAPI-stained fluorescence images of AGS cells treated with STO-609 for indicated time points (8, 24, 48, and 72 h), showing reduced cell density and increased multinucleated cells (highlighted), indicative of defects in cell cycle progression and mitotic regulation. All experiments were performed in biological triplicate. Data are presented as mean ± SEM. Statistical significance was determined using Student’s *t*-test.

These findings establish the functional relevance of CAMKK2 signaling in maintaining the malignant properties of gastric cancer cells and provide a strong biological rationale for subsequent phosphoproteomic and kinase-centric analyses aimed at defining the signaling mechanisms underlying these phenotypic effects.

To systematically characterize the kinome and signaling alterations downstream of CAMKK2, we performed a global phosphoproteomic analysis following pharmacological inhibition of CAMKK2 in gastric cancer cells. AGS cells were treated with STO-609, a selective CAMKK2 inhibitor, and subjected to TMT-based quantitative phosphoproteomic profiling, followed by integrative bioinformatics and functional analyses. An overview of the experimental design is shown in Figure 1.

Based on our previous dose–response analyses, STO-609 was used at its IC₅₀ concentration (18.5 µM), as determined by MTT assays in AGS cells. This experimental strategy enabled unbiased interrogation of CAMKK2-dependent phosphorylation networks and downstream signaling pathways in gastric cancer.

### Quantitative phosphoproteomic analysis upon inhibition of CAMKK2 in gastric cancer cells

Given the phenotypic alterations observed upon CAMKK2 inhibition in gastric cancer (GC) cells, we next sought to characterize the global phosphorylation changes underlying these effects. To this end, we employed a TMT-based quantitative phosphoproteomics approach to systematically investigate alterations in phosphorylation signaling in the AGS gastric cancer cell line following CAMKK2 inhibition. A schematic overview of the experimental workflow is shown in Figure 2.

**Figure 2.**
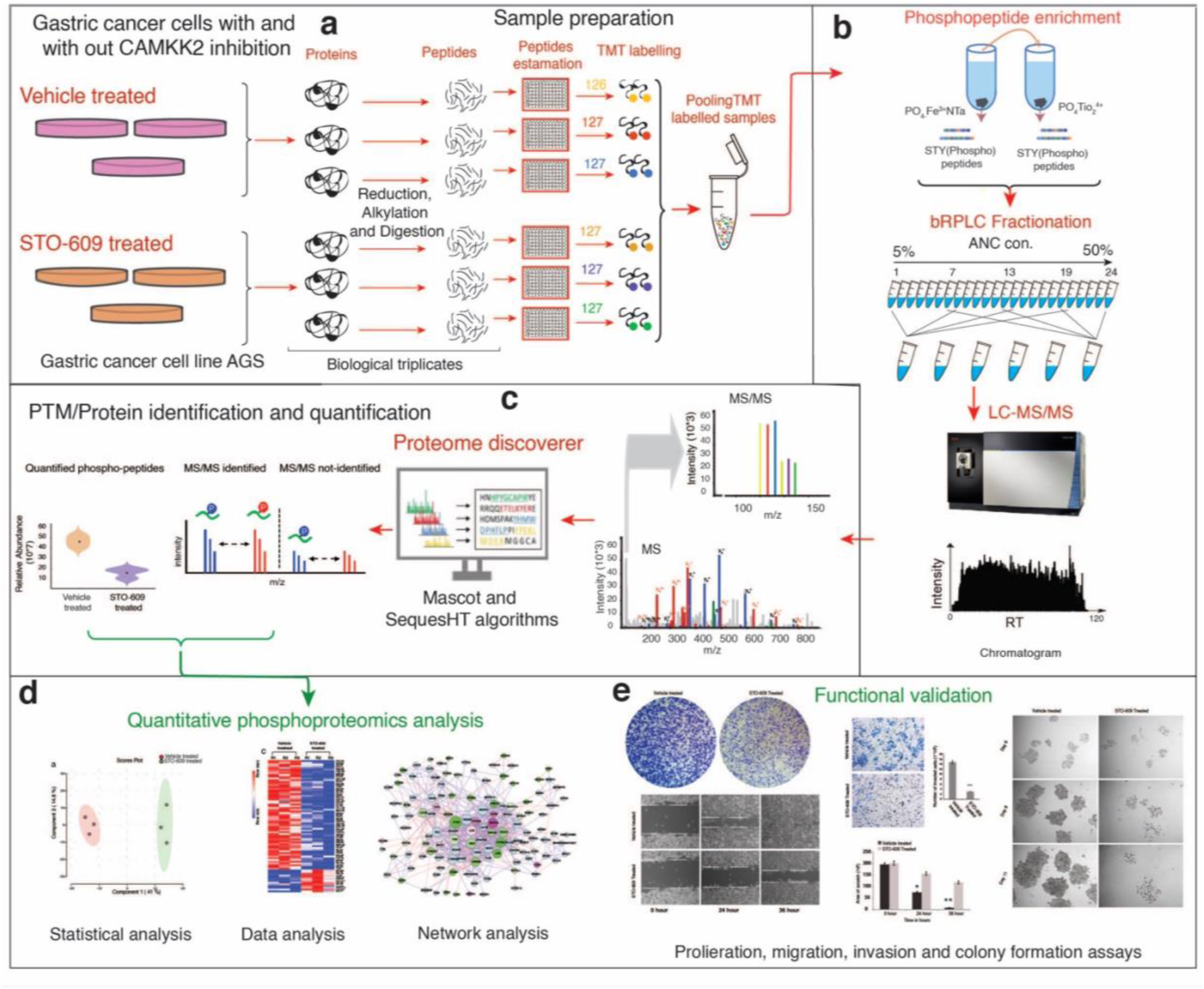
Experimental workflow for quantitative phosphoproteomic analysis of CAMKK2 signaling in gastric cancer: **(a)** AGS gastric cancer cells were treated with the CAMKK2 inhibitor STO-609 or vehicle control in biological triplicates, followed by protein extraction, tryptic digestion, peptide quantification, and TMT labeling. Labeled peptides were pooled for downstream analysis. **(b)** Pooled peptides were subjected to phosphopeptide enrichment using Fe³⁺-NTA and TiO₂ affinity-based methods, followed by basic reverse-phase liquid chromatography (bRPLC) fractionation. **(c)** Enriched phosphopeptides were analyzed by LC-MS/MS using an Orbitrap Fusion Tribrid mass spectrometer, and data were processed using Proteome Discoverer with Mascot and SequestHT algorithms. **(d)** Quantitative phosphoproteomic and bioinformatics analyses were performed to identify differentially phosphorylated proteins, signaling pathways, and kinase networks regulated by CAMKK2. **(e)** Functional validation of CAMKK2-dependent signaling was carried out using proliferation, colony formation, migration, and invasion assays.

Our phosphoproteomic analysis resulted in the identification and quantification of 10,523 phosphopeptides, corresponding to 3,120 proteins and 9,603 unique phosphosites, at a false discovery rate (FDR) of 1%. The overall distribution of phosphopeptide abundances across control and CAMKK2-inhibited samples demonstrated high consistency among biological replicates (Figure 3A).

**Figure 3.**
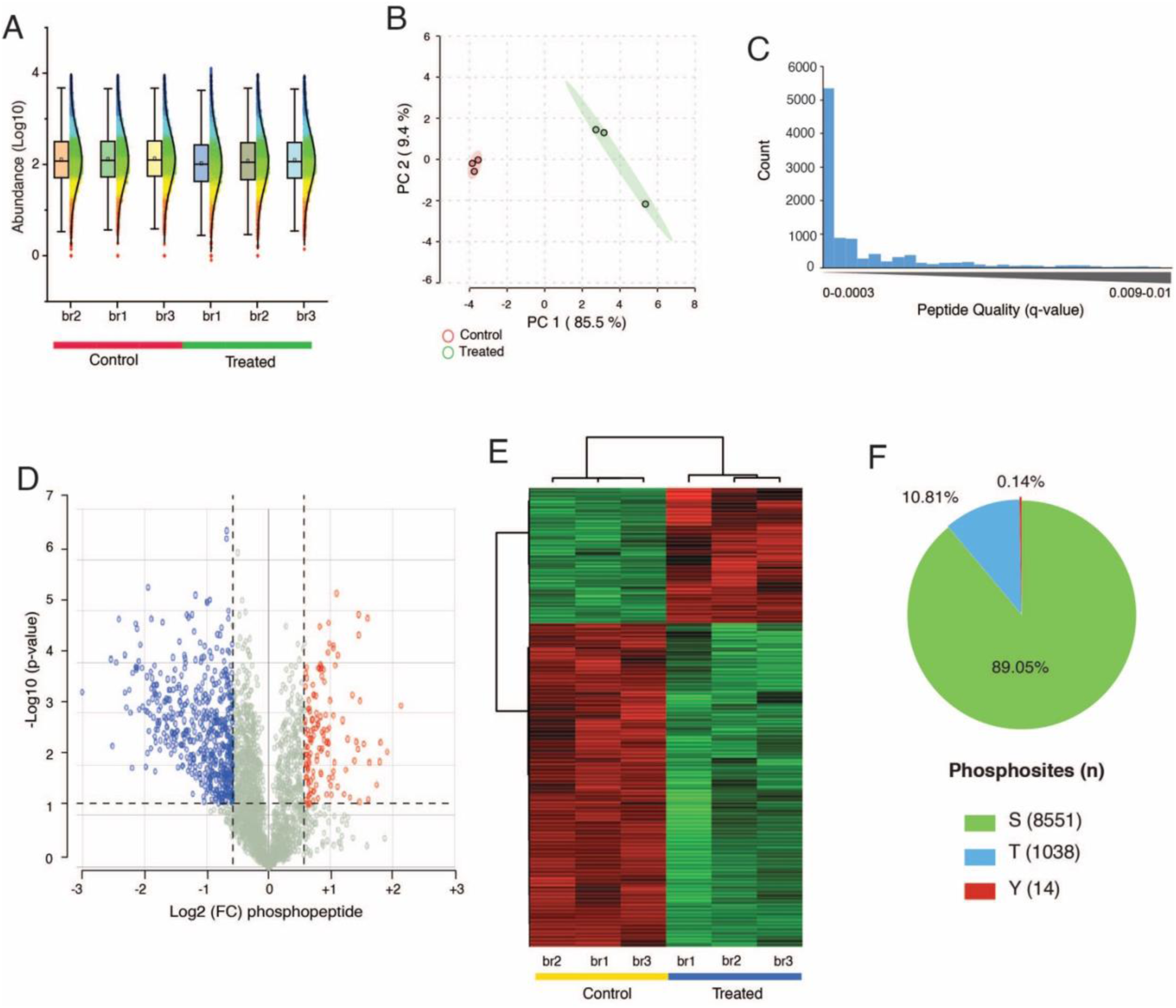
Quantitative phosphoproteomic profiling reveals extensive phosphorylation changes upon CAMKK2 inhibition: **(A)** Box and violin plots showing the distribution of log₁₀-transformed phosphopeptide abundances across biological replicates of vehicle treated and STO-609 treated AGS cells, demonstrating high reproducibility among samples. **(B)** Principal component analysis (PCA) of phosphoproteomic data showing clear separation between control and CAMKK2-inhibited samples, indicating distinct phosphorylation profiles upon CAMKK2 inhibition. **(C)** Distribution of phosphopeptide identification confidence based on q-values, demonstrating that the majority of identified phosphopeptides exhibit high confidence (low q-values). **(D)** Volcano plot depicting differentially phosphorylated phosphopeptides upon CAMKK2 inhibition. Significantly hypophosphorylated and hyperphosphorylated phosphopeptides are highlighted based on fold-change and statistical significance thresholds. **(E)** Heatmap of significantly altered phosphopeptides showing hierarchical clustering of control and STO-609 treated samples, illustrating coordinated phosphoproteomic remodeling following CAMKK2 inhibition. **(F)** Pie chart showing the distribution of identified phosphorylation sites across serine (S), threonine (T), and tyrosine (Y) residues.

To assess data quality, we examined phosphopeptide q-values, which reflect the confidence of peptide-spectrum matches in LC-MS/MS analyses. The majority of identified phosphopeptides exhibited q-values below 0.001, indicating high confidence in peptide identification and overall dataset robustness (Figure 3B).

To further evaluate reproducibility and treatment-specific differences, principal component analysis (PCA) was performed. PCA revealed tight clustering among biological replicates and clear separation between control and CAMKK2-inhibited samples, demonstrating strong experimental reproducibility and distinct phosphorylation profiles upon CAMKK2 inhibition (Figure 3C).

Differential phosphorylation analysis identified 1,101 significantly hypophosphorylated phosphopeptides (≥1.5-fold change), corresponding to 752 proteins, and 301 significantly hyperphosphorylated phosphopeptides (≥1.5-fold change), corresponding to 234 proteins, following CAMKK2 inhibition in AGS cells. A comprehensive list of all identified and differentially regulated phosphopeptides is provided in Tables S2 and S3. The global distribution of phosphorylation changes is visualized using a volcano plot, highlighting significantly altered phosphopeptides upon CAMKK2 inhibition (Figure 3D).

Hierarchical clustering analysis of significantly altered phosphopeptides revealed distinct phosphorylation patterns between control and treated samples, further confirming robust and coordinated phosphoproteomic remodeling following CAMKK2 inhibition (Figure 3E).

Finally, analysis of phosphosite composition revealed that the majority of identified phosphorylation events occurred on serine residues (89.05%), followed by threonine residues (10.81%), with a small fraction on tyrosine residues (0.14%), consistent with the known distribution of phosphorylation sites in eukaryotic cells (Figure 3F).

Collectively, these results demonstrate that CAMKK2 inhibition induces extensive and reproducible alterations in the phosphoproteome of gastric cancer cells, providing a high-quality foundation for downstream functional, pathway, and kinase-centric analyses.

### Functional analysis of the CAMKK2-regulated phosphoproteome

Given the widespread signaling alterations observed upon CAMKK2 inhibition, we next performed comprehensive bioinformatics analyses to functionally annotate the differentially phosphorylated proteins. Enrichment analyses were carried out to categorize these proteins based on cellular localization, molecular function, biological process, signaling pathways, transcription factor associations, and microRNA networks. Gene Ontology (GO) enrichment analysis was performed using the g:Profiler web-based platform.

Cellular component analysis revealed that the majority of CAMKK2-regulated phosphoproteins were predominantly localized to the nucleus, followed by the nucleoplasm and nuclear lumen, indicating that CAMKK2 inhibition primarily impacts nuclear signaling events (Figure 3, Cellular Component).

Molecular function analysis demonstrated that CAMKK2-regulated phosphoproteins were significantly enriched in functions related to protein binding, RNA binding, histone binding, and cell adhesion molecule binding, highlighting the involvement of CAMKK2 in regulating protein protein and protein RNA interaction networks (Figure 4, Molecular Function).

**Figure 4.**
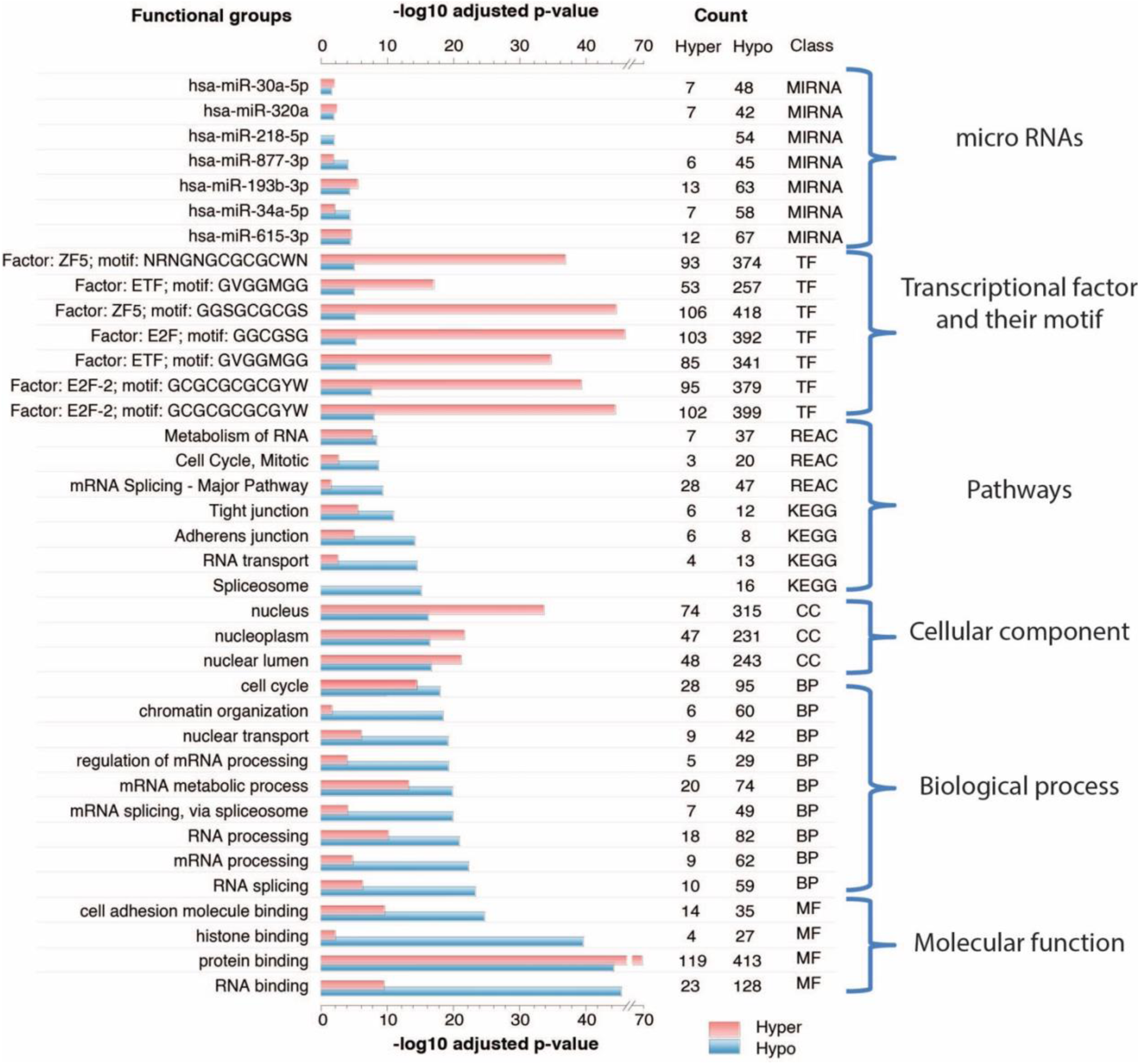
Functional enrichment analysis of CAMKK2-regulated phosphoproteome: Bar plot showing functional enrichment of differentially phosphorylated proteins following CAMKK2 inhibition in AGS gastric cancer cells. Gene Ontology (GO), pathway, transcription factor, and microRNA enrichment analyses were performed using the g:Profiler platform. Enriched categories are grouped into microRNAs, transcription factors and their DNA-binding motifs, signaling pathways, cellular components, biological processes, and molecular functions, as indicated. Bars represent log₁₀ adjusted *p*-values for significantly enriched terms, with red and blue bars denoting hyperphosphorylated and hypophosphorylated protein sets, respectively. The number of hyper and hypophosphorylated proteins contributing to each category is indicated.

Consistent with these observations, biological process enrichment analysis showed that differentially phosphorylated proteins upon CAMKK2 inhibition were involved in a broad spectrum of nuclear and regulatory processes, including cell cycle regulation, mRNA metabolic processes, RNA processing, RNA splicing, and nuclear transport (Figure 4, Biological Process). Pathway enrichment analysis further identified multiple signaling pathways affected by CAMKK2 inhibition in gastric cancer. Notably, a substantial proportion of hypophosphorylated proteins were associated with pathways involved in RNA metabolism, mRNA splicing, spliceosome assembly, and RNA transport. In addition, hypophosphorylated proteins were also enriched in pathways related to cell cycle progression, mitosis, tight junctions, and adherens junctions, suggesting that CAMKK2-dependent phosphorylation is critical for maintaining both proliferative signaling and cellular structural integrity (Figure 4, Pathways).

To further investigate transcriptional regulatory mechanisms associated with CAMKK2 inhibition, transcription factor enrichment analysis was performed using the TRANSFAC database through the g:Profiler platform. This analysis revealed significant enrichment of transcription factors, including ZF5, ETF, E2F, and E2F-2, indicating that CAMKK2-regulated phosphorylation events converge on transcriptional programs governing cell cycle control and gene expression (Figure 4, Transcription Factors).

Finally, microRNA enrichment analysis was conducted using the miRTarBase database to explore potential post-transcriptional regulatory networks associated with the altered phosphoproteome. This analysis identified several microRNAs linked to CAMKK2 inhibition, including hsa-miR-30a-5p, hsa-miR-193b-3p, and hsa-miR-615-3p, suggesting that CAMKK2 signaling may indirectly modulate gene expression through miRNA-associated regulatory pathways (Figure 4, microRNAs).

Collectively, these analyses demonstrate that CAMKK2 inhibition induces extensive remodeling of the phosphoproteome, with pronounced effects on nuclear signaling, RNA metabolism, cell cycle regulation, and transcriptional and post-transcriptional control, providing mechanistic insight into the role of CAMKK2 in gastric cancer signaling networks.

### Kinase substrate enrichment analysis reveals CAMKK2-dependent regulation of cell cycle and MAPK signaling networks

To identify upstream kinases responsible for the observed phosphorylation changes following CAMKK2 inhibition, we performed kinase substrate enrichment analysis (KSEA) using the differentially phosphorylated phosphosites. This analysis enabled systematic inference of kinase activity changes based on the enrichment patterns of known kinase substrates.

Heatmap based visualization of kinase substrate interactions revealed extensive rewiring of phosphorylation networks upon CAMKK2 inhibition (Figure 5A-B). Multiple kinases displayed significant enrichment across differentially phosphorylated proteins, indicating coordinated regulation of kinase-driven signaling pathways rather than isolated phosphorylation events.

**Figure 5.**
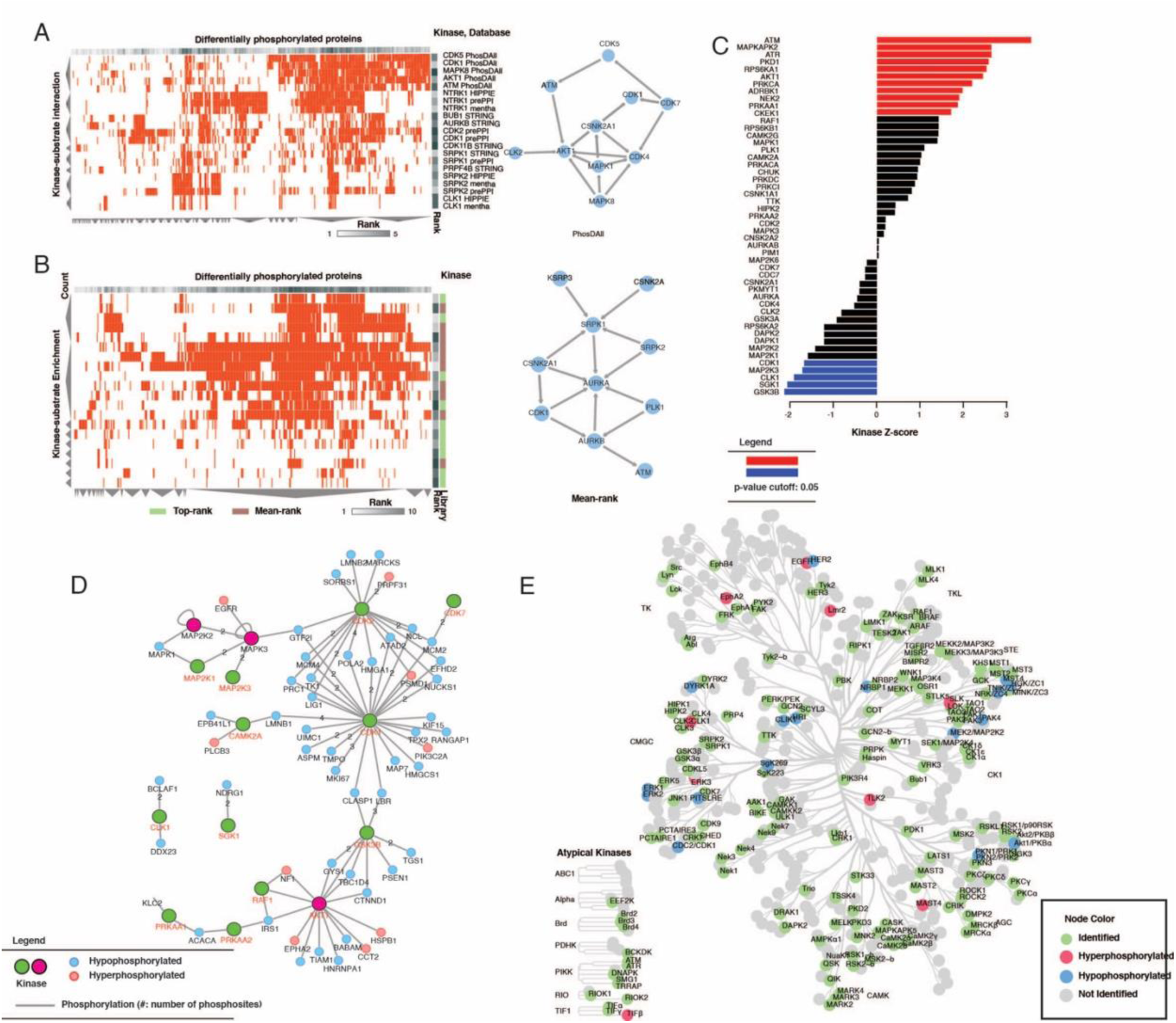
Kinase substrate enrichment analysis reveals CAMKK2-dependent kinase signaling alterations: **(A)** Heatmap showing enrichment of kinase substrate interactions among differentially phosphorylated proteins following CAMKK2 inhibition, with network visualization highlighting key kinases within the inferred signaling architecture. **(B)** Mean-rank–based kinase–substrate enrichment heatmap and corresponding network representation identifying kinases consistently regulated upon CAMKK2 inhibition. **(C)** Kinase activity changes represented as Z-scores derived from substrate phosphorylation patterns, highlighting kinases with significantly altered activity (p ≤ 0.05). **(D)** Kinase substrate interaction network integrating phosphorylation status of substrates, where node colors indicate hyper- or hypophosphorylation and edge thickness reflects the number of phosphosites. **(E)** Global kinome map illustrating the distribution of identified and differentially phosphorylated kinases across major kinase families following CAMKK2 inhibition.

Network-based representation of kinase substrate relationships highlighted key kinases occupying central positions within the inferred signaling architecture (Figure 5A-B, right panels). Notably, kinases associated with cell cycle control, DNA damage response, and stress signaling, including CDK family members, MAPKs, ATM, and CK2, emerged as prominent nodes, suggesting that CAMKK2 activity is functionally linked to these regulatory axes.

To quantitatively assess kinase activity changes, kinase Z-scores were calculated based on substrate phosphorylation patterns. This analysis identified a subset of kinases with significantly altered activity following CAMKK2 inhibition (Figure 5C). Among the most positively enriched kinases were MAPKAPK2, ATM, ATR, PRKDC, AKT1, and MAPK family members, whereas kinases associated with mitotic progression and glycogen metabolism exhibited reduced activity. These findings indicate that CAMKK2 inhibition leads to broad modulation of kinase signaling programs governing proliferation, stress responses, and survival.

To further contextualize these findings, we constructed a kinase–substrate interaction network integrating phosphorylation status and connectivity (Figure 5D). This analysis revealed that several kinases, including CAMKK2-associated signaling nodes, were connected to both hyper-and hypophosphorylated substrates, underscoring their role as integrative regulators of phosphorylation dynamics. Importantly, many substrates within this network were involved in DNA replication, chromatin organization, RNA processing, and cytoskeletal regulation, consistent with the functional enrichment patterns observed in Figure 3.

Finally, global kinome mapping was performed to visualize the distribution of identified kinases across established kinase families (Figure 5E). This analysis demonstrated that CAMKK2 inhibition affected kinases spanning multiple families, including CMGC, AGC, CAMK, TK, and atypical kinases, with a notable enrichment of kinases involved in cell cycle regulation and MAPK signaling. Both hyper and hypophosphorylated kinases were distributed across these families, highlighting the extensive remodeling of kinase signaling networks upon CAMKK2 inhibition.

Collectively, these analyses demonstrate that CAMKK2 inhibition leads to widespread reprogramming of kinase activity, impacting key signaling pathways that regulate cell cycle progression, MAPK signaling, DNA damage response, and RNA metabolism. These results position CAMKK2 as a central upstream regulator coordinating multiple kinase-driven signaling networks in gastric cancer.

### Sequence motif analysis reveals loss of CDK/MAPK-associated phosphorylation signatures upon CAMKK2 inhibition

To further investigate whether the phosphorylation changes observed upon CAMKK2 inhibition reflected specific kinase recognition patterns, we performed sequence motif analysis of differentially phosphorylated phosphosites using MoMo (The MEME Suite, https://meme-suite.org/meme/tools/momo). Motifs were analyzed separately for phosphosites associated with CAMKK2 dependent and MAPK dependent signaling to delineate kinase-specific sequence preferences.

Sequence logo analysis of hypophosphorylated CAMKK2-associated substrates revealed enrichment of serine and proline-directed motifs, characterized by basic residues upstream of the phosphorylation site and proline enrichment at the +1 position (Figure 6A). These sequence features are consistent with consensus motifs recognized by cell cycle associated kinases, including CDK and related CMGC family members, suggesting attenuation of proliferative kinase signaling upon CAMKK2 inhibition.

**Figure 6.**
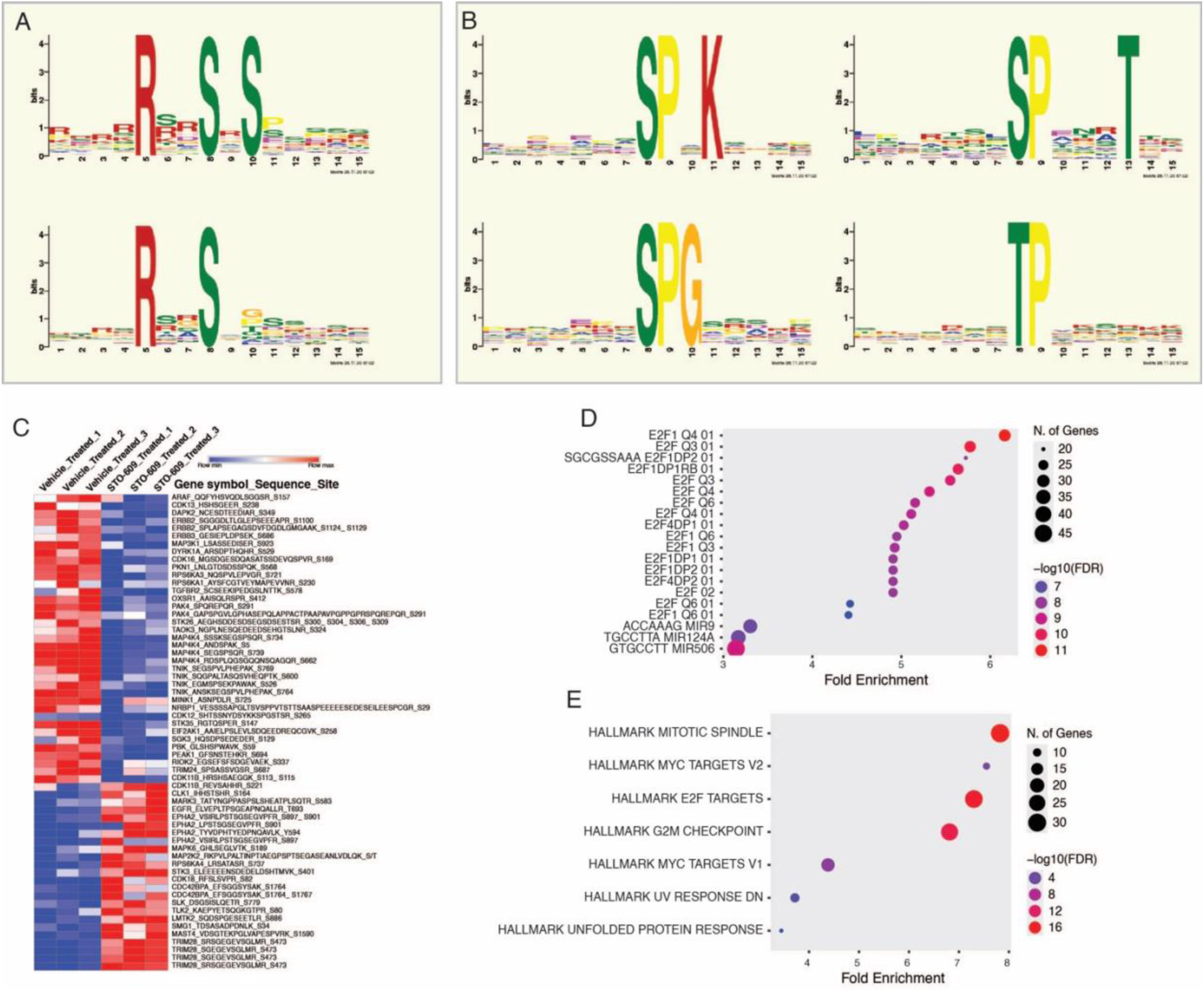
Sequence motif and pathway enrichment analyses reveal loss of CDK/MAPK-associated phosphorylation programs upon CAMKK2 inhibition: **(A)** Sequence logo representations of enriched phosphorylation motifs among hypophosphorylated CAMKK2-associated substrates, highlighting serine-centered and proline-directed consensus motifs. **(B)** Sequence logo representations of enriched phosphorylation motifs among hypophosphorylated MAPK-associated substrates, showing characteristic SP and TP motifs indicative of MAPK activity. **(C)** Heatmap depicting phosphorylation patterns of kinase substrates across vehicle-treated and STO-609 treated AGS cells, illustrating coordinated reduction in phosphorylation following CAMKK2 inhibition. **(D)** Motif enrichment analysis showing over-representation of E2F transcription factor associated motifs among hypophosphorylated proteins, with dot size representing the number of genes and color indicating statistical significance (−log₁₀ FDR). **(E)** Hallmark gene set enrichment analysis of hypophosphorylated proteins highlighting significant enrichment of pathways related to mitotic spindle assembly, G2/M checkpoint regulation, E2F targets, and MYC-driven transcriptional programs.

Similarly, motif analysis of MAPK associated hypophosphorylated substrates demonstrated prominent enrichment of SP and TP motifs, a hallmark of MAPK and proline directed kinase activity (Figure 6B). The presence of these canonical MAPK recognition sequences supports the kinase substrate enrichment results shown in Figure 4 and indicates reduced MAPK mediated phosphorylation following CAMKK2 inhibition.

To examine the consistency of these motif-associated phosphorylation changes across biological replicates, we next visualized phosphorylation patterns of kinase substrates using a heatmap representation (Figure 6C). This analysis revealed a clear separation between vehicle-treated and CAMKK2-inhibited samples, with widespread reduction in phosphorylation of kinase substrates involved in cell cycle regulation, chromatin organization, and signal transduction, underscoring the robustness and reproducibility of the observed phosphorylation changes.

Given the strong association of identified motifs with cell cycle related kinases, we next assessed whether motif containing proteins were linked to specific transcriptional regulatory programs. Motif enrichment analysis using MSigDB revealed significant over-representation of E2F transcription factor associated motifs among hypophosphorylated proteins (Figure 6D). Multiple E2F motif variants, including E2F1 and E2F-DP associated signatures, were enriched, indicating convergence of CAMKK2 dependent phosphorylation events on E2F regulated gene programs. Consistent with this observation, Hallmark gene set enrichment analysis further demonstrated significant enrichment of pathways associated with mitotic spindle assembly, G2/M checkpoint regulation, E2F targets, and MYC driven transcriptional programs among hypophosphorylated proteins (Figure 6E). These findings suggest that CAMKK2 dependent phosphorylation preferentially supports core proliferative and mitotic programs, and that inhibition of CAMKK2 results in coordinated suppression of these fundamental cellular processes.

Collectively, these motif-based analyses demonstrate that CAMKK2 inhibition leads to selective loss of phosphorylation at CDK- and MAPK-associated consensus motifs, converging on E2F driven cell cycle and mitotic programs. These results provide sequence level mechanistic support for the kinase centric and functional pathway alterations observed in Figures 3 and 4.

### Proposed model of CAMKK2-dependent regulation of kinase signaling in gastric cancer

Based on integrated phenotypic, phosphoproteomic, kinase-centric, and motif-based analyses, we propose a model in which CAMKK2 functions as a central upstream regulator of proliferative kinase signaling in gastric cancer (Figure 7). Pharmacological inhibition of CAMKK2 using STO-609 leads to coordinated attenuation of downstream kinase networks, including CDK-, MAPK-, and mitotic kinase–associated signaling, resulting in widespread hypophosphorylation of proteins involved in cell cycle progression, RNA processing, and cytoskeletal organization. This signaling collapse converges on suppression of E2F-driven transcriptional programs, disruption of mitotic processes, and impaired Rho GTPase–mediated cytoskeletal dynamics, ultimately manifesting as reduced proliferation, clonogenic growth, migration, and invasion of gastric cancer cells. Collectively, these findings position CAMKK2 as a key signaling hub that integrates multiple kinase-driven pathways to sustain malignant phenotypes in gastric cancer and highlight CAMKK2 inhibition as a potential therapeutic strategy for targeting aberrant proliferative signaling.

**Figure 7.**
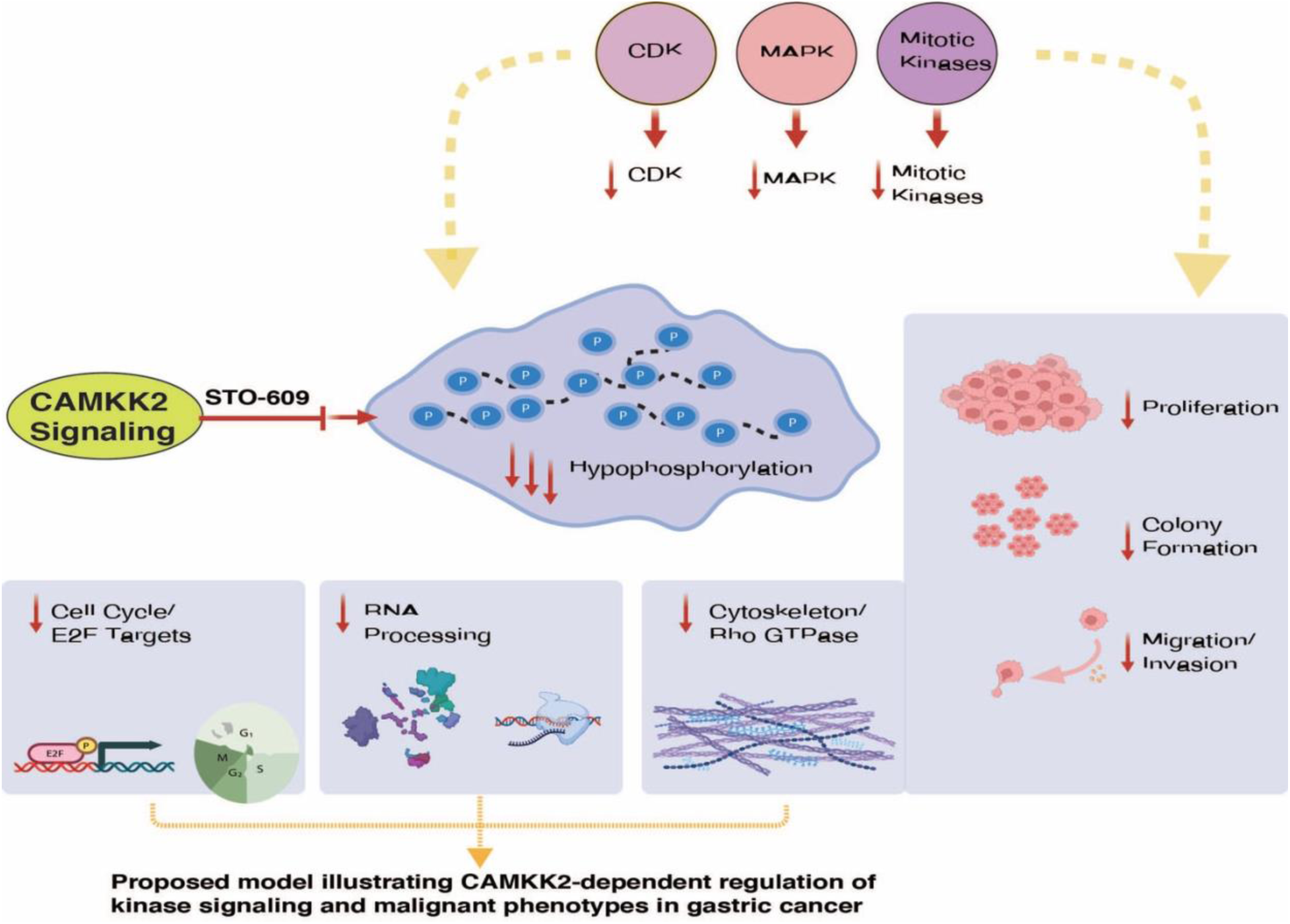
Proposed model of CAMKK2-dependent regulation of kinase signaling in gastric cancer: Schematic model illustrating CAMKK2 as a central regulator of kinase signaling in gastric cancer. Inhibition of CAMKK2 with STO-609 attenuates CDK, MAPK, and mitotic kinase-associated signaling, leading to widespread hypophosphorylation of downstream substrates. These changes suppress E2F-driven cell-cycle programs, disrupt RNA processing and cytoskeletal dynamics, and ultimately reduce proliferation, clonogenic growth, migration, and invasion in gastric cancer cells.

## Discussion

Calcium/calmodulin-dependent protein kinase kinase 2 (CAMKK2) has been reported to be overexpressed in multiple cancer types, including gastric cancer, and has been implicated in the regulation of oncogenic signaling pathways that promote cellular proliferation and survival [8, 17, 24, 25]. Analyses of public cancer datasets, including TCGA, along with prior experimental studies, have demonstrated elevated CAMKK2 expression at both the transcript and protein levels in gastric tumors [8]. Despite these observations, the signaling mechanisms by which CAMKK2 contributes to gastric tumorigenesis have remained incompletely characterized, particularly at the level of global kinase signaling and phosphorylation network regulation.

Protein kinases and their dysregulation represent central drivers of cancer progression, and targeting kinase signaling has been a major focus of cancer therapeutics for several decades [26–28]. Advances in mass spectrometry-based phosphoproteomics have enabled comprehensive profiling of kinase activity and downstream signaling events, providing unprecedented insight into oncogenic signaling networks [29–31]. In the present study, we applied a quantitative phosphoproteomic approach to systematically characterize CAMKK2-dependent signaling in gastric cancer cells, integrating phenotypic assays with global phosphorylation, functional enrichment, kinase-centric, and motif-based analyses.

Consistent with previous reports linking CAMKK2 to oncogenic growth, pharmacological inhibition of CAMKK2 using STO-609 resulted in marked suppression of gastric cancer cell proliferation and clonogenic potential [17]. In addition, CAMKK2 inhibition significantly impaired migratory and invasive behavior, indicating a broader role for CAMKK2 in regulating malignant phenotypes beyond cell growth alone. The emergence of multinucleated cells upon prolonged CAMKK2 inhibition further suggests disruption of cell cycle progression and mitotic fidelity, a phenotype frequently associated with deregulated kinase signaling in cancer.

Mitogen-activated protein kinase (MAPK) signaling pathways are evolutionarily conserved kinase modules that link extracellular stimuli to fundamental cellular processes such as proliferation, differentiation, migration, and apoptosis [32–35]. Dysregulation of the RAS/RAF/MEK/ERK cascade is a hallmark of many solid tumors, including gastric cancer, where aberrant MAPK signaling contributes to tumor progression and therapeutic resistance [36–38]. While MAPK activation has traditionally been attributed to upstream receptor tyrosine kinases, Ras, or RAF mutations, alternative regulatory inputs sustaining MAPK signaling in gastric cancer are less well defined [38–40]. Our kinase substrate enrichment and activity inference analyses reveal that CAMKK2 inhibition leads to coordinated attenuation of MAPK-associated phosphorylation programs, suggesting that CAMKK2 contributes to the maintenance of MAPK signaling output in gastric cancer cells.

In addition to MAPK signaling, our data indicate that CAMKK2 inhibition impacts cyclin-dependent kinases (CDKs) and mitotic kinases, which are central regulators of cell-cycle progression. CDKs orchestrate orderly cell cycle transitions and are frequently deregulated in cancer through aberrant activation or loss of regulatory control [41–43]. Previous studies have demonstrated extensive crosstalk between MAPK signaling and CDK activation, with MAPK-driven pathways promoting CDK activity and cell cycle progression [38, 44, 45]. In this study, reduced phosphorylation of CDK-associated substrates, together with enrichment of cell-cycle–related biological processes and transcriptional programs, suggests that CAMKK2-dependent signaling supports proliferative CDK activity in gastric cancer cells.

Minichromosome maintenance (MCM) proteins and other DNA replication factors are critical downstream targets of CDK-mediated phosphorylation and play essential roles in DNA replication licensing and S-phase progression [46, 47]. Aberrant phosphorylation of replication-associated proteins has been widely reported in cancer and is associated with uncontrolled proliferation [48, 49]. Our phosphoproteomic analysis revealed reduced phosphorylation of multiple nuclear and chromatin-associated proteins involved in DNA replication and RNA processing following CAMKK2 inhibition, indicating that CAMKK2-dependent kinase signaling supports replication competence and transcriptional regulation in gastric cancer.

Beyond nuclear and cell cycle–associated processes, CAMKK2 inhibition also resulted in significant alterations in cytoskeletal organization and Rho GTPase-associated signaling pathways. Rho GTPases and cytoskeletal regulators play critical roles in cell shape, motility, and invasion, and their dysregulation is closely linked to cancer metastasis [50, 51]. The impaired migration and invasion observed upon CAMKK2 inhibition are therefore consistent with the phosphoproteomic evidence indicating suppression of cytoskeletal and adhesion-related phosphorylation programs, suggesting that CAMKK2 integrates signaling pathways controlling both proliferative and invasive phenotypes.

Sequence motif analysis provided further mechanistic insight into the kinase specificity of CAMKK2-regulated phosphorylation events. Hypophosphorylated sites were enriched for proline-directed motifs characteristic of CDK- and MAPK-mediated phosphorylation, as well as motifs associated with mitotic kinase activity. Importantly, motif-based enrichment analyses converged on E2F transcription factor-associated programs, which are central regulators of cell cycle progression and DNA replication. Dysregulation of E2F signaling has been extensively linked to genomic instability, mitotic defects, and tumorigenesis [52–54], providing a plausible mechanistic link between CAMKK2 inhibition, loss of kinase signaling, and the multinucleated phenotype observed in this study.

Collectively, these findings support a model in which CAMKK2 functions as an upstream signaling node that coordinates multiple kinase-driven pathways, including MAPK, CDK, and mitotic kinase networks, to sustain malignant phenotypes in gastric cancer. Inhibition of CAMKK2 leads to widespread hypophosphorylation of proteins involved in cell cycle regulation, RNA metabolism, and cytoskeletal dynamics, resulting in suppression of proliferative, clonogenic, migratory, and invasive behaviors. While kinase activity was inferred from substrate phosphorylation patterns and further validation will be required, this systems-level analysis provides a comprehensive view of CAMKK2-dependent signaling architecture in gastric cancer.

In summary, by integrating phenotypic characterization with global phosphoproteomics, kinase-centric analyses, and sequence motif enrichment, our study advances current understanding of CAMKK2 signaling in gastric cancer. These findings highlight CAMKK2 as a central regulator of oncogenic kinase networks and suggest that targeting CAMKK2 may represent a promising therapeutic strategy for disrupting aberrant proliferative and invasive signaling in gastric cancer.

## Conclusion

In conclusion, this study provides a comprehensive systems-level view of CAMKK2-dependent phosphorylation signaling in gastric cancer. Through integrated phenotypic assays, quantitative phosphoproteomics, kinase-substrate enrichment, and motif analyses, we demonstrate that CAMKK2 functions as an upstream regulatory hub coordinating multiple kinase-driven signaling networks, including CDK, MAPK, and mitotic kinase-associated pathways. Inhibition of CAMKK2 results in widespread hypophosphorylation of proteins involved in cell cycle progression, transcriptional regulation, RNA processing, and cytoskeletal organization, leading to impaired proliferative and invasive phenotypes. These findings extend previous pathway focused observations by revealing the broader kinase signaling architecture downstream of CAMKK2 and establish a mechanistic link between CAMKK2 activity, global phosphorylation remodeling, and malignant behavior in gastric cancer cells. Collectively, our work highlights CAMKK2 as a critical integrator of oncogenic kinase signaling and supports its potential as a therapeutic target for disrupting aberrant proliferative signaling in gastric cancer.

## Ethics approval and consent to participate

Not applicable

## Data Availability Statement

Data related to the study are provided in the manuscript and supplementary rest of the data and can be provided request.

## Competing Interests

The authors declare no relevant financial or non-financial interests.

## Funding

No funding.

## Author Contributions

P. K. M. conceived the idea, designed the experiments and interpretation of data, and critically reviewed and edited the manuscript. M.A.N conceived the idea, designed the experiments along with P.K.M. performed experiments and data analysis, drafted the manuscript, and prepared figures. T.S.K.P. reviewed and edited the manuscript. All authors read and approved the final version of the manuscript. All the authors agree to publish the manuscript.

## Acknowledgements

The authors gratefully acknowledge Yenepoya (Deemed to be University) for providing the infrastructure and state-of-the-art mass spectrometry facility required to conduct this study exclusively within our institution. We also acknowledge the support provided by the Department of Biotechnology (DBT) through the National Facility grant under the project “Skill Development in Mass Spectrometry-based Metabolomics Technology BIC” (BT/PR40202/BTIS/137/53/2023).

## Abbreviations

CAMKK2: Calcium/calmodulin-dependent protein kinase kinase
GC: Gastric cancer
TMT: Tandem mass tag
LC-MS/MS: Liquid chromatography tandem mass spectrometry
GO: Gene Ontology
KSEA: Kinase substrate enrichment analysis
MAPK: Mitogen activated protein kinase
CDK: Cyclin-dependent kinase
ERK: Extracellular signal-regulated kinase
AKT: Protein kinase B
ATM: Ataxia telangiectasia mutated
ATR: Ataxia telangiectasia and Rad3-related
PRKDC: Protein kinase DNA-activated catalytic subunit
E2F: E2 factor transcription factor
PCA: Principal component analysis
FDR: False discovery rate
PTM: Post-translational modification
Fe-NTA: Ferric nitrilotriacetic acid
TiO₂: Titanium dioxide
ACN: Acetonitrile
TFA: Trifluoroacetic acid
TEABC: Triethylammonium bicarbonate
SEM: Standard error of the mean
SDS-PAGE: Sodium dodecyl sulfate–polyacrylamide gel electrophoresis
DHB: 2,5-Dihydroxybenzoic acid
X2K: eXpression2Kinases.

## Notes

### Competing Interest Statement

The authors have declared no competing interest.

